# Dengue virus 2 capsid protein chaperones strand displacement without altering the capsid-coding region hairpin element’s structural functionality

**DOI:** 10.1101/2021.01.17.427046

**Authors:** Xin Ee Yong, Palur Venkata Raghuvamsi, Ganesh S. Anand, Thorsten Wohland, Kamal K. Sharma

## Abstract

By virtue of its chaperone activity, the capsid protein of dengue virus strain 2 (DENV2C) promotes nucleic acid structural rearrangements. However, the role of DENV2C during the interaction of RNA elements involved in stabilizing the 5’-3’ panhandle structure of DENV RNA is still unclear. Therefore, we determined how DENV2C affects structural functionality of the capsid-coding region hairpin element (cHP) during RNA rearrangement of the 9-nt conserved sequence (5CS) to its complementary 3CS counterpart. The cHP element has two distinct functions: a role in translation start codon selection and a role in RNA synthesis. Our results showed that the cHP hairpin impedes annealing between the 5CS and the 3CS elements. Although DENV2C does not modulate structural functionality of the cHP hairpin, it accelerates annealing and specifically promotes strand displacement of 3CS during 5’-3’ panhandle formation. Furthermore, DENV2C exerts its chaperone activity by favoring one of the active conformations of the cHP element. Based on our results, we propose mechanisms for annealing and strand displacement involving the cHP element. Thus, our results provide mechanistic insights on how DENV2C regulates RNA synthesis by modulating essential RNA elements in the capsid-coding region, that in turn allow for DENV replication.

## INTRODUCTION

Like all flaviviruses, dengue virus (DENV) is an enveloped, non-polyadenylated positive-strand RNA virus with a type 1 cap structure located at the 5’ end of the genome and 5’ and 3’ untranslated regions (UTRs) flanking a single open reading frame (ORF) (1). DENV causes dengue fever and its more severe and potentially lethal manifestations, dengue hemorrhagic fever/dengue shock syndrome. The four serotypes of DENV (DENV1–4) are transmitted to humans via two mosquito species: *Aedes aegypti* or *Aedes albopictus* and are the leading cause of arboviral diseases worldwide (2, 3). The DENV2 replication cycle begins when the virus binds and enters a susceptible host cell by receptor-mediated endocytosis, leading to the release of the viral genome into the cytoplasm. The DENV2 ORF is subsequently translated as a polyprotein from a start codon located at the 5’ end of the region encoding the viral capsid protein (DENV2C). The polyprotein is cleaved into three structural (envelope [E], pre-membrane [prM] and capsid [DENVC)] and seven non-structural proteins (NS1, NS2A, NS2B, NS3, NS4A, NS4B, NS5). The genomic RNA (vRNA) is synthesized by the viral replicase complex via a negative-sense intermediate, and newly transcribed vRNA undergo late rounds of translation that are dependent upon RNA synthesis.

The 5’UTR of DENV2 contains several conserved cis-acting RNA elements that play a key role in regulating viral translation and RNA synthesis (3–6). The 5’ conserved sequence (5CS) regulates RNA synthesis through interaction with the complementary 3CS, located within the 3’UTR (7, 8) (Figure 1A). The downstream AUG region (DAR) was identified, with the 5DAR located just downstream of the initiating AUG and the complementary 3DAR sequence found in the 3’UTR (9, 10). The 5DAR/3DAR acts together with the 5CS/3CS and 5’/3’ upstream AUG region (5UAR/3UAR) to circularize the vRNA (Figure 1A), which is a requirement for the efficient RNA synthesis but is not involved in viral translation (5, 8–11).

**Figure 1:**
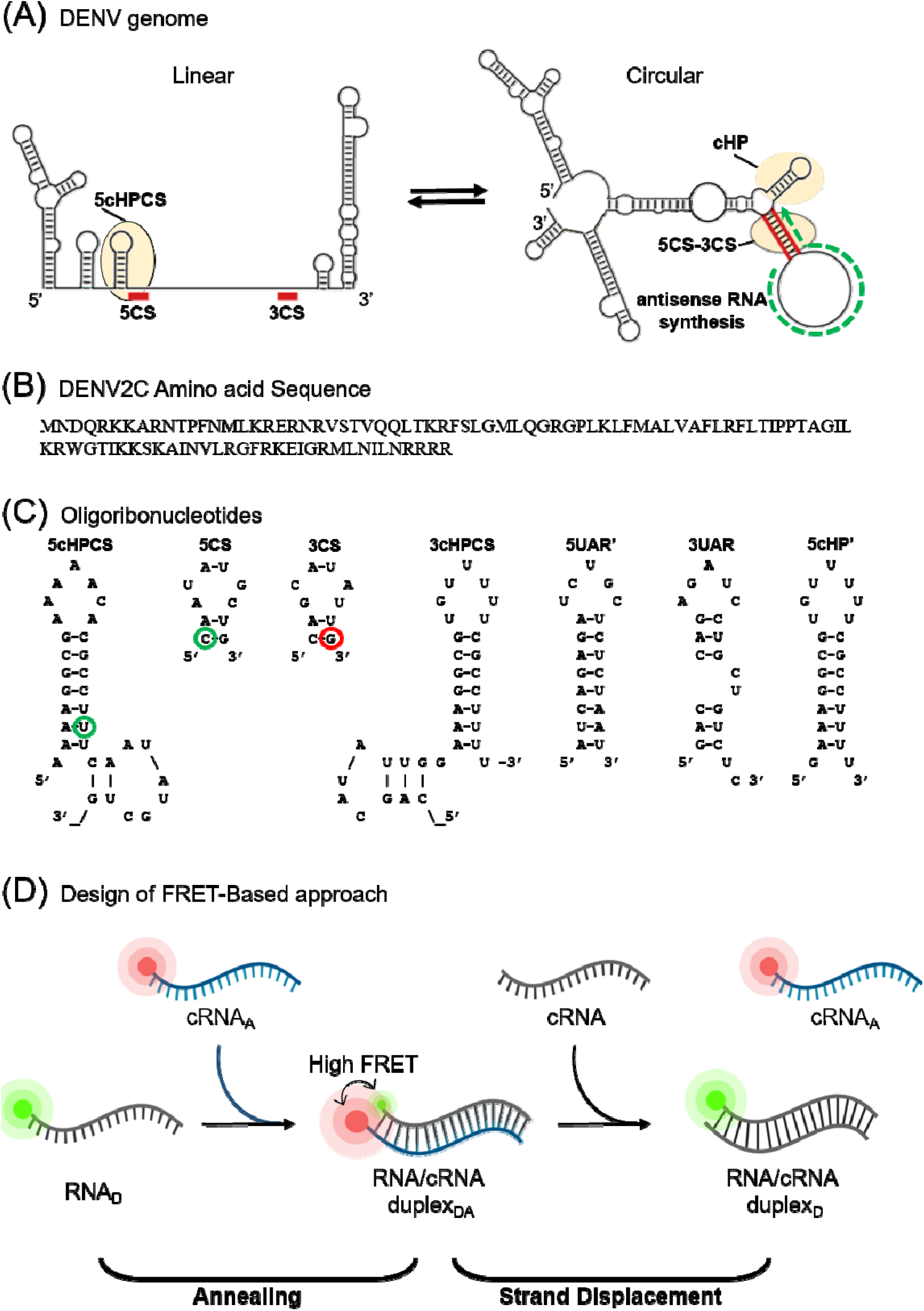
Schematic representation of DENV genome, DENV2C, ORN sequences and FRET-based annealing and strand displacement approach used in this study. **(A) Interplay between two conformations of DENV genome during 5’-3’ panhandle formation.** The 5’ and 3’ UTR region of DENV genome contain conserved sequences, 5CS and 3CS (shown by red blocks). The 5CS is downstream of the cHP region (shown by yellow region) and is considered a sequence-independent essential hairpin structure. Annealing (shown by red lines) of these conserved sequences (5CS/3CS) allow the formation of the circularized genome. On the other hand, the synthesis of antisense DENV RNA (shown by green arrow) leads to strand displacement between the 5CS and 3CS sequences. **(B) DENV2C amino acid sequence.** The DENV2C protein is a homodimer with both basic and acidic residues. In the protein sequence of DENV2C monomer, the first 21 residues constitute the N-terminus disordered region. **(C) Oligoribonucleotides.** ORN sequences are derived from the 5’UTR and 3’UTR regions of the genome. Their secondary structures were predicted using the mfold webtool (http://unafold.rna.albany.edu/). 5cHPCS and 5CS are labelled with FAM as donor at the 21^st^ position and 5’ end, respectively (shown by green circles) while 3CS is labelled with TAMRA as acceptor at 3’ end (shown by red circle). **(D) Schematic representation of the FRET-Based approach.** The annealing of RNA_D_-cRNA_A_ will lead to the formation of a duplex that is labelled with both FAM (green) and TAMRA (red). Due to the close proximity of donor and acceptor dyes, a high FRET efficiency and a decrease in the FAM intensity is observed. The addition of cRNA during the strand displacement reaction led to the displacement of cRNA_A_, which shifts the two dyes apart from each other, resulting in a decrease in FRET efficiency and the restoring of FAM fluorescence.

Interestingly, a cis-acting RNA secondary structure, termed the capsid-coding hairpin (cHP) (Figure 1A), was found to regulate both viral translation and RNA synthesis in a sequence-independent manner (12, 13). Intriguingly, how the cHP influences DENV vRNA synthesis is still unknown. The cHP element briefly stalls the scanning initiation complex over the first AUG, favouring its recognition (13) in the absence of a strong initiation context. When 5CS and 3CS have annealed, cHP has a stretched-out stem structure that includes the first nucleotide (nt) of 5CS (14) (5cHPCS in Figure 1A). Once the vRNA has switched from its role as an mRNA to a template for vRNA synthesis, cyclization decreases the length of the cHP stem slightly in the presence of the 3’UTR (14) (shown by 5cHPCS sequence in Figure 1A). We hypothesize that the cHP element stabilizes the overall 5⍰-3⍰ panhandle structure (12, 13) of DENV vRNA during RNA synthesis, probably via an interplay between annealing and unpairing/strand displacement of the 5CS and 3CS elements (Figure 1A). Consequently, the cHP and the CS elements, in association with the other mentioned cyclization motifs (DAR and UAR), could regulate DENV vRNA in adopting different conformations via both annealing as well as strand displacement events (9). This interplay of DENV vRNA from circular to linear conformations modulates negative and positive-strand RNA synthesis (15).

In addition, it is believed that proteins like NS5 and NS3 helicase assist this interplay of DENV vRNA by unwinding secondary/tertiary structures and displacing the double-stranded (ds) regions in vRNA (16, 17). We have shown that DENV2C modulates the annealing and melting of the RNA elements in 5’UTR region of DENV2 vRNA and plays a role in genomic rearrangements (18). DENV2C is an RNA chaperone, shown to promote the annealing of the hammerhead ribozyme to its substrate as well as the dissociation of the cleaved substrates (19). As an RNA chaperone, DENV2C helps to prevent the misfolding of RNA by allowing RNA to sample many different conformations (20). DENV2C exists as a homodimer, consisting of two 100-aa subunits (Figure 1B). The protein is highly basic and has a flexible disordered region in the N-terminus, both key characteristics of many RNA chaperones (21–24). The protein associates with the vRNA to form the nucleocapsid and the nucleocapsid is encapsulated in a lipid bilayer containing E and M proteins (25). DENV2C binds to vRNA via its basic residues in helix-4 and binds to the lipid bilayer using its hydrophobic cleft in helix-1 (26, 27). Considering the chaperone activity of DENV2C, we hypothesize that this protein plays a significant role both in annealing and strand displacement of the 5CS and 3CS elements during stabilization of the 5’-3’ panhandle structure by modulating the cHP element (12, 13).

Since the cHP regulates RNA synthesis in a sequence-independent manner (12), we investigated the structural functionality of the cHP element during the annealing and strand displacement kinetics of the 9-nt 5CS to its complementary 3CS (Figure 1C). We monitored annealing kinetics using either 32-nt 5cHPCS or 9-nt 5CS, labelled with 6-carboxyfluorescein (FAM) as the donor fluorophore (Figure 1C), to the complementary 3CS labelled with carboxytetramethylrhodamine (TAMRA) as the acceptor (Figure 1C), forming a Förster resonance energy transfer (FRET)-pair. With the addition of the acceptor-labelled complementary 3CS to donor-labelled 5cHPCS or 5CS, the proximity between donor and acceptor dyes increases and will lead to the quenching of the donor fluorescence during real-time annealing kinetics (Figure 1D). Likewise, we monitored strand displacement of the acceptor-labelled 3CS from these annealed 5cHPCS/3CS or 5CS/3CS duplexes by adding excess concentrations of non-labelled 3CS that will lead to fluorescence recovery of the donor (Figure 1D). By monitoring the annealing kinetics of 5cHPCS and 5CS with 3CS, we found that the presence of the cHP hairpin structure impedes the annealing between 5CS and 3CS. However, no strand displacement was observed in both annealing complexes. Interestingly, DENV2C accelerates both 5cHPCS/3CS and 5CS/3CS annealing and promotes strand displacement of 3CS from both annealed duplexes. Moreover, we observed that DENV2C specifically nucleates the strand displacement of 3CS without disturbing the structural functionality of the cHP hairpin. We also determined, using time-resolved FRET (trFRET), that DENV2C exerts its chaperoning activity by favoring one of the active conformations of 5cHPCS/3CS and 5CS/3CS duplexes. We further proposed mechanisms for the 5cHPCS/3CS and 5CS/3CS annealing as well as DENV2C-promoted 3CS-strand displacement from these annealed complexes using a coupled annealing and strand displacement approach (Figure 1D). Overall, our results suggest a reaction mechanism where the cHP structure stabilizes the 5’-3’ panhandle. In addition, we also determined that DENV2C modulates the role of the cHP during the 5CS-3CS annealing as well as during strand displacement. Therefore, our results improve the understanding of *cis*-acting RNA elements and reveal how DENV2C regulates RNA synthesis through the stabilization and genomic rearrangements of the essential RNA elements in the capsid-coding region that regulate DENV fitness.

## MATERIALS AND METHODS

### Oligoribonucleotides (ORN)

All ORNs were synthesized by Integrated DNA Technologies (Singapore). The donor-labelled 5cHPCS and 5CS were synthesized with 6-carboxyfluorescein (FAM) at the 21^st^ position and the 5’ end, respectively (Figure 1C). The acceptor-labelled 3CS were synthesized with carboxytetramethylrhodamine (TAM) at the 3’ end (Figure 1C). All ORNs were purified by the manufacturer using HPLC.

### DENV2C protein synthesis and purification

DENV2C protein was expressed and purified as described in an earlier study (18). DENV2C protein preparations were ribonuclease-free (18) and RNA-free, determined by a A260/A280 value of 0.7 for the purified protein, very close to the theoretical value of 0.57 for a protein sample not contaminated by nucleic acids.

### Fluorescence spectroscopy

Fluorescence spectroscopy measurements were done on a Cary Eclipse Fluorescence Spectrophotometer (Agilent) with a temperature control module, using Hellma^®^ fluorescence cuvettes with an internal chamber volume of 45 μL. Excitation and emission wavelengths of 480 nm and 520 nm were used to track the intensity of FAM in real-time. Annealing reactions were performed in second order conditions by adding acceptor-labelled ORN to donor-labelled ORN at a 1:1 ratio while the strand displacement reactions were performed in pseudo first order conditions with the concentration of non-labelled ORN being at least 10-fold more than the doubly-labelled annealed complexes. Equal volumes of both reactants were mixed at the start of the reaction to prevent high local concentrations of either reactant. A final concentration of 2 μM DENV2C was used for monitoring the effect of DENV2C either on annealing or on strand displacement reactions. All reactions were performed in 50 mM HEPES, 30 mM NaCl, 0.2 mM MgCl_2_, pH 7.5 buffer and at 20°C. Reactions could not be monitored at higher temperatures (>20°C) due to rapid and non-reproducible reactions. All curve fitting was done on OriginProTM software (ver 9.55).

### Time-resolved FRET (trFRET)

trFRET measurements were carried out on a commercial Olympus FV1200 laser scanning confocal microscope equipped with a time-resolved LSM upgrade kit (Microtime 200, PicoQuant, GmbH, Berlin, Germany). Donor-labelled ORNs was excited with a 485 nm pulsed diode laser with a 20 MHz repetition rate and 29 mW power (PDL series, Sepia II combiner module). The beam was focused into the sample by a water immersion objective (60×, NA 1.2; Olympus, Singapore) after being reflected by a dichroic mirror (DM405/485/543/635, Olympus, Singapore) and the scanning unit. The fluorescence was collected by the same objective followed by a pinhole (120 mm) to remove out-of-focus light. The fluorescence signal was spectrally divided into donor (green) and acceptor (red) channels by a 560 DCLP mirror. The FAM donor fluorescence was recorded by a SPAD (SPCM-AQR-14, PerkinElmer Optoelectronics, Quebec, Canada), through a 513/17 band pass emission filter (Omega, VT). This donor signal was further processed by a time correlated single photon counting card (TimeHarp 260, PicoQuant) to build up the histogram of photon arrival times. The trFRET measurements were recorded for 180 s at 20°C. The mean lifetime (τ) was calculated from the individual fluorescence lifetimes (τ_i_) and their relative amplitudes (a_i_) according to (τ) = ∑*α*_*i*_*τ*_*i*_ Donor fluorescence lifetime decay data were treated using the software SymPhoTime 64 (PicoQuant, GmbH). In all cases, the *χ*^2^ values were close to 1 and the weighted residuals as well as their autocorrelation were distributed randomly around 0, indicating a good fit. The reported values are mean and S.D.’s from at least three replicates.

## RESULTS

### 5CS/3CS annealing is 2-folds faster in the absence of cHP hairpin

We monitored annealing kinetics of the donor-labelled 5cHPCS to its CS-region-complementary and acceptor-labelled 3CS by tracking the decrease in FAM fluorescence (Figure. 2A). Comparing the emission spectra of the donor-labelled 5cHPCS at the start and end of the annealing reaction shows an increase in FRET with the formation of the duplex that contains both FAM and TAMRA in close proximity as seen by a decrease in FAM fluorescence (Figure S1). The real-time fluorescence intensity traces of the 5cHPCS/3CS reaction kinetics were fitted to a monoexponential equation [1], where *I*(*t*) is the fluorescence intensity at 520 nm, upon excitation at 480 nm, *k*_obs_ is the second order reaction rate and *t*_0_ is the start time of the reaction. *I*_*0*_ and *I*_*f*_ is the fluorescence intensity of the donor-labelled 5cHPCS where the 5CS region is in single-strand state (in absence of acceptor-labelled 3CS) and in the double-strand duplex form due to presence of the acceptor-labelled 3CS, respectively.

[1]

**Figure 2:**
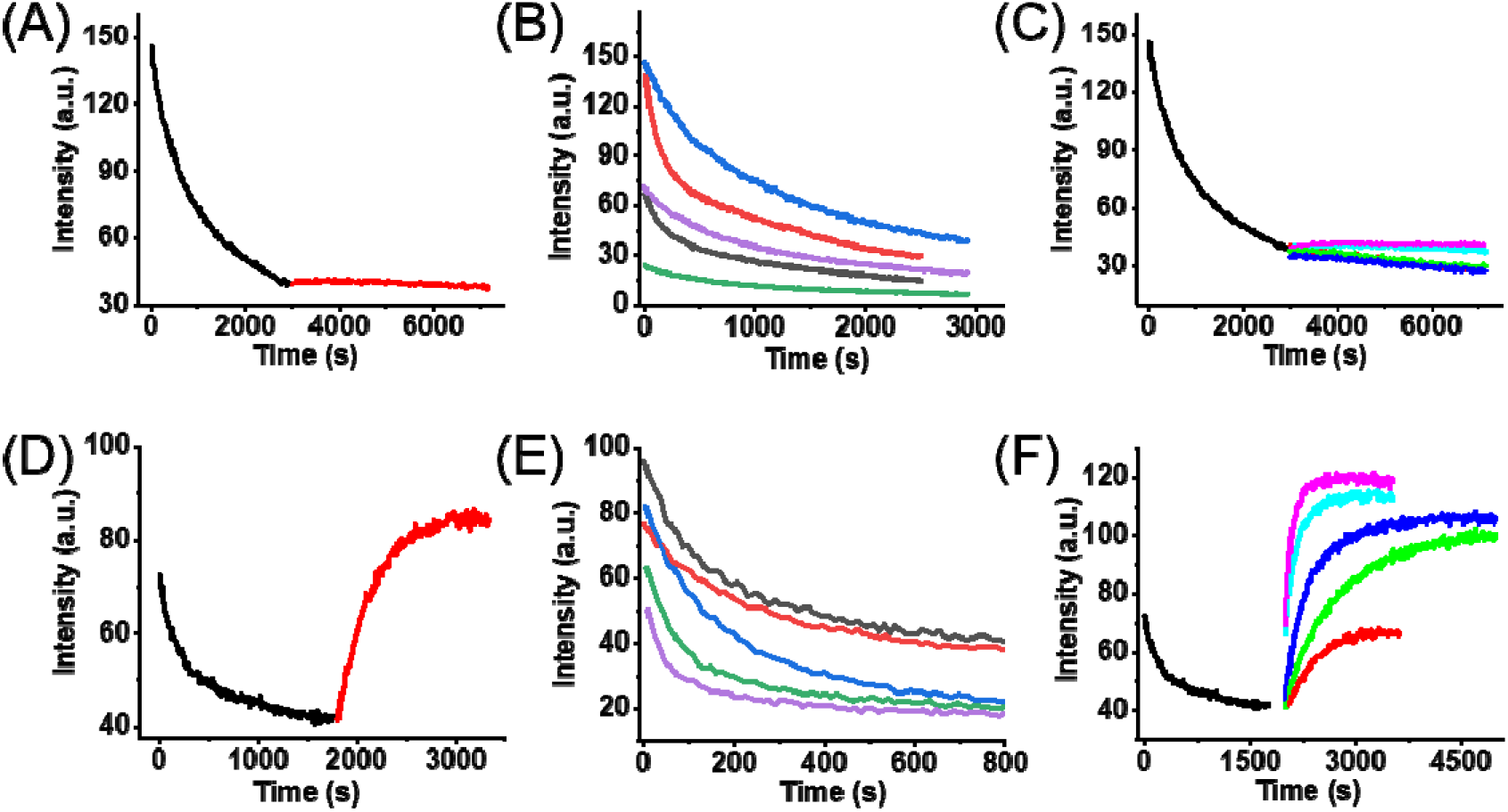
Real-time progress curves of 5cHPCS/3CS annealing and 5cHPCS/3CS’ strand displacement in the absence (A-C) and presence (D-F) of DENV2C. Progress curves of 10nM FAM-labelled 5cHPCS annealing (black traces) with 10nM TAMRA-labelled 3CS and strand displacement with 300 nM non labelled 3CS in the (A) absence and (D) presence of DENV2C. Progress curves of 10nM FAM-labelled 5cHPCS/3CS annealing with increasing concentration of TAMRA-labelled [3CS] in (B) absence (blue-20nM, red-50 nM, pink-70 nM, dark grey-100 nM and green-150 nM) and (E) presence (dark grey-10 nM, red-20 nM, blue-30 nM, green-50 nM and pink-100 nM) of 2 μM DENV2C. Progress curves of 5cHPCS/3CS’ strand displacement initiating from the duplex formed during 5cHPCS/3CS annealing in (C) absence (red-100 nM, green-500 nM, blue-1 μM, cyan-1.5 μM and pink-2 μM) and (F) presence (red-100 nM, green-500 nM, blue-1 μM, cyan-1.5 μM and pink-2 μM) of DENV2C. All annealing and strand displacement traces were fitted using equation [1] and the obtained values are provided in Figure 3. Excitation and emission wavelengths used were 480 nm and 520 nm, respectively.

The reaction rates, *k*_obs_, were plotted against concentrations of the acceptor-labelled 3CS and was observed to be linearly varying with increasing 3CS concentrations ([3CS)] (Figure 3A and 2B). The linear relationship between *k*_obs_ and [3CS] follows equation [2] (27–29).

[2]

**Figure 3:**
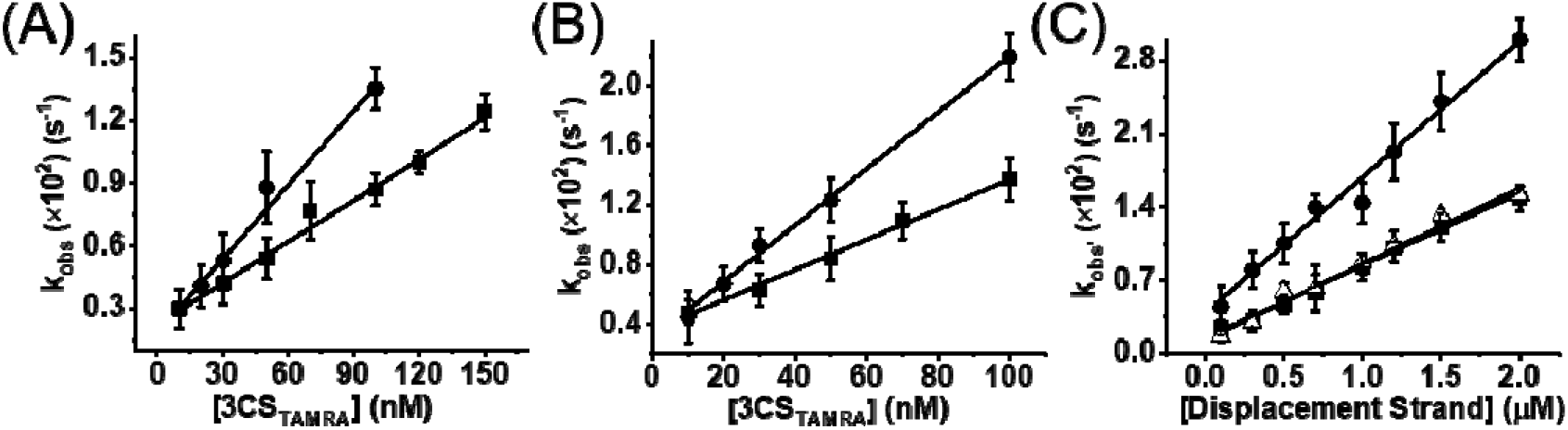
Kinetic parameters of (A) 5cHPCS/3CS and (B) 5CS/3CS annealing and (C) 5cHPCS/3CS’, 5CS/3CS’ and 5cHPCS/3cHPCS strand displacement in the absence and presence of DENV2C. Kinetic parameters were obtained by fitting real-time progress curves shown in figure 2. During (A) 5cHPCS/3CS and (B) 5CS/3CS annealing, the obtained values of apparent rates (*k*_obs_) were plotted against increasing concentrations of the complementary TAMRA-labelled 3CS in (squares) absence and (circles) presence of DENV2C. (C) Kinetic rates (*k*_obs’_) for the displacement of TAMRA-labelled 3CS during 5cHPCS/3CS’ (squares), 5CS/3CS’ (circles) and 5cHPCS/3cHPCS (open triangles) were also plotted against increasing concentration of displacement strands (3CS’ and 3cHPCS). All plotted kinetic rates were fitted to equation [2] and the obtained values are provided in Table 1. Excitation and emission wavelengths used were 480 nm and 520 nm, respectively. Error bars show standard deviation from at least three repeats.

Fitting the linear plot of *k*_obs_ against increasing [3CS] with equation [2] generated the kinetic parameters shown in Table 1. Based on the acquired kinetic parameters and a continuous decrease in donor intensity plateau with increasing [3CS] (Figure 2B), a reaction mechanism with single kinetic pathway can be proposed:

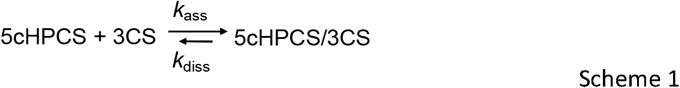

where the final stable extended duplex, 5cHPCS/3CS, is formed through a bimolecular reaction (29, 30). The formation of the 5cHPCS/3CS duplex is governed by the second order association constant, *k*_ass_, and the first order dissociation constant, *k*_diss_, respectively. We observed the *k*_ass_ value of 66.3 (± 3) × 10^3^ M^−1^s^−1^ and *k*_*diss*_ value of 23.4 (± 2.8) × 10^−4^ s^−1^ for the 5cHPCS/3CS annealing (Table 1). Since the value of k_ass_ is at least 2 orders of magnitude smaller than the rate constants reported for annealing of unstructured sequences (10^5^ −10^7^ M^−1^s^−1^), it seems that the 5CS region of the 5cHPCS undergoes structural rearrangement during the formation of the extended duplex in the presence of the complementary 3CS sequence. We validate the postulated annealing mechanism (31–34) using the Dynafit numerical resolution software (35), which allows simultaneous fitting of the experimental progress curves obtained at different [3CS] (Figure S2). The estimated elementary rate constants *k*_ass_ and *k*_diss_ (Table S1) were in excellent agreement with those found by the empirical approach (Table 1) and validated the proposed annealing mechanism for the 5cHPCS/3CS reaction.

**Table 1:**
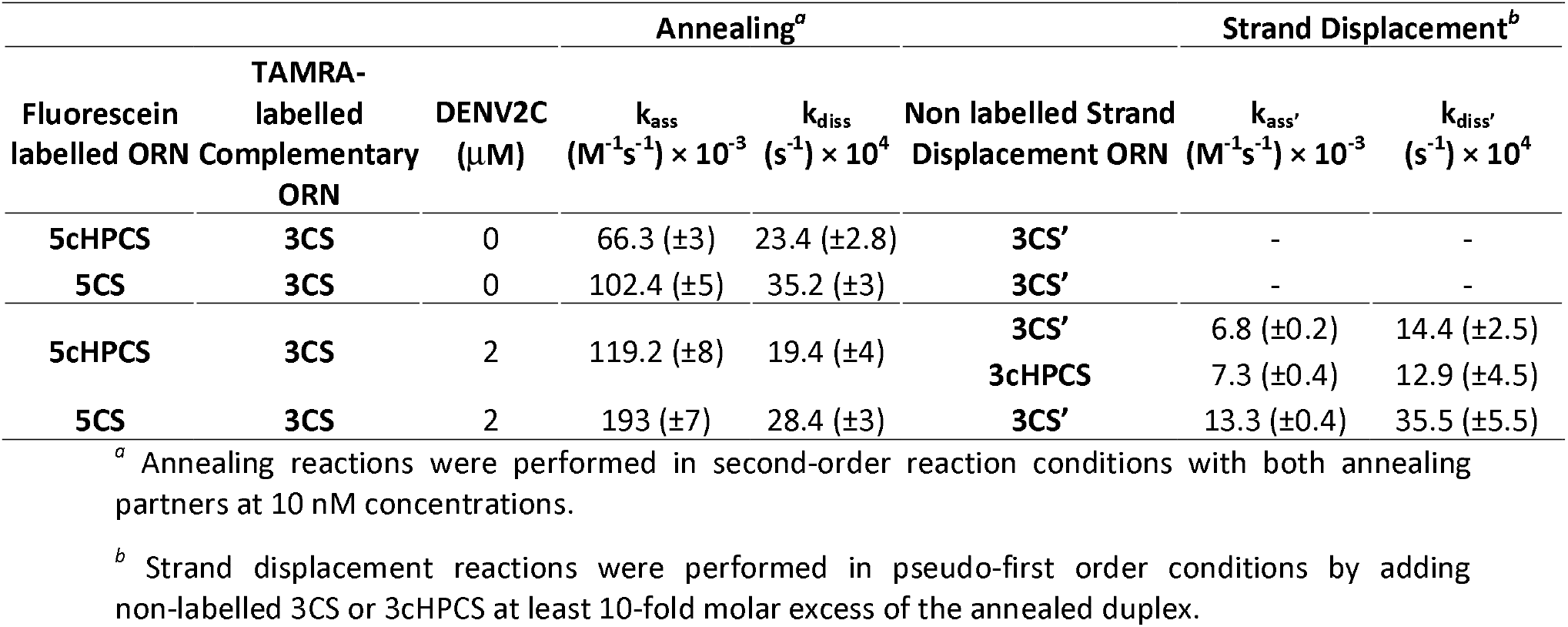
Kinetic parameters of 5cHPCS/3CS and 5CS/3CS annealing, and 5cHPCS/3CS’, 5CS/3CS’ and 5cHPCS/3cHPCS strand displacement in the absence and presence of DENV2C. Kinetic rate constants, *k*_ass_, *k*_diss_, *k*_ass’_ and *k*_diss’_, were calculated from the dependence of the *k*_obs_ values on the concentration of the unlabelled ORN, using equation [2] as indicated in Figure 3.

Similar to 5cHPCS/3CS annealing, 5CS/3CS annealing shows a linear dependence of the *k*_obs_ values (Figure 3B) as well as a continuous decrease in donor intensity plateau against increasing [3CS], indicating a similarity of reaction mechanism to that proposed by Scheme 1 (shown by Scheme a in the supplementary). The 5CS/3CS annealing reaction was found to be ~2-fold faster as compared to the 5cHPCS/3CS annealing reaction (by comparing corresponding *k*_ass_ values in Table 1), indicating a hindrance in 5cHPCS/3CS annealing due to the presence of the cHP hairpin structure. The cHP hairpin RNA element serves only a structural function in 5cHPCS/3CS annealing, and yet it seems to modulate the annealing of 5CS to its complementary 3CS. On the other hand, similar values of *k*_*diss*_ for both 5cHPCS/3CS and 5CS/3CS annealing suggest that the cHP region has limited effect on the stability of 5CS-3CS duplex formation (Table 1).

### DENV2C accelerates both 5cHPCS/3CS and 5CS/3CS annealing

Since DENV2C has been shown to chaperone annealing of essential RNA elements (UAR and cHP) from the DENV2 5’UTR region (18), we investigated if DENV2 chaperones the annealing of the 5cHPCS/3CS and 5CS/3CS and whether it alters the hindering effect of the cHP element. We mimicked large molecular ribonucleoprotein (RNP) complex conditions (36–38) by using DENV2C at a DENV2C:ORN molar ratio of 100:1 to characterize the annealing of 5cHPCS/3CS and 5CS/3CS (Figure S3A). Addition of DENV2C to the donor-labelled 5cHPCS and 5CS sequences led to a decrease in their fluorescence (compare black emission spectra in Figure S1A and S1B, respectively), indicating the condensation of RNA sequences due to the binding of DENV2C with ORNs in RNP complex conditions (36–38). Annealing of 5cHPCS/3CS and of 5CS/3CS were then triggered by adding acceptor-labelled 3CS to the preformed complex of either 5cHPCS or 5CS and DENV2C (Figure 2D). The real-time fluorescence intensity traces of both the DENV2C-promoted 5cHPCS/3CS (Figure 2D) and 5CS/3CS (Figure S3C) reaction kinetics showed monoexponential decrease in FAM intensity and were fitted with equation [1]. Interestingly, both DENV2C-promoted 5cHPCS/3CS and 5CS/3CS annealing showed linear dependence of the *k*_obs_ values as well as a continuous decrease in donor intensity plateau against increasing [3CS] (Figure 3A and 3B) and thus a similar reaction mechanism to that proposed by Scheme 1 can be suggested. As expected, DENV2C accelerated both 5cHPCS/3CS and 5CS/3CS annealing, and a ~2-fold increase in *k*_obs_ in the presence of DENV2C was observed when compared to the corresponding annealing in the absence of the protein (Table 1). Incidentally, DENV2C-promoted 5CS/3CS annealing was still found to be ~2-fold faster as compared to DENV2C-promoted 5cHPCS/3CS annealing (by comparing the corresponding *k*_*ass*_ values in Table 1), suggesting that DENV2C does not regulate the structural functionality of cHP hairpin in the 5cHPCS/3CS annealing.

### DENV2C promotes strand displacement of 3CS from both 5cHPCS/3CS and 5CS/3CS complexes

Next, we monitored strand displacement kinetics of the acceptor-labelled 3CS from preformed donor/acceptor-labelled 5cHPCS/3CS and 5CS/3CS duplexes by adding non-labelled 3CS in at least 10-fold molar excess. Due to pseudo-first order conditions, the addition of non-labelled 3CS sequences should displace acceptor-labelled 3CS from either the 5cHPCS/3CS or the 5CS/3CS duplex leading to an increase and opposite decrease in the FAM and TAMRA fluorescence, respectively. Contrary, no changes in the strand displacement kinetic traces and the emission spectra of the dual-labelled 5cHPCS/3CS (Figure 2C and S1A) or 5CS/3CS (Figure S3B) duplexes at the start and end of the strand displacement reaction were observed, thus suggesting that no strand displacement takes place in these conditions. We tested up to 200-fold excess non-labelled 3CS to investigate strand-displacement but could not detect any increase in FAM fluorescence suggesting the stability of both 5cHPCS/3CS or 5CS/3CS annealed duplexes (Figure 2C and S3B). Intriguingly, addition of non-labelled 3CS, even at 10-fold excess, leads to a rapid increase in FAM fluorescence in the presence of DENV2C (Figure 2D and S4B). The result proves the chaperoning role of the DENV2C in strand displacement of the acceptor-labelled 3CS and thus aiding the genomic recombination during DENV RNA synthesis.

The real-time fluorescence intensity traces of DENV2C-promoted 3CS strand displacement from either 5cHPCS/3CS (Figure 2F) or 5CS/3CS (Figure S3C) annealed duplex were fitted to equation [1]. *I(t)* is the actual fluorescence intensity at 520 nm, upon excitation at 480 nm, *I’*_*0*_ and *I’*_*f*_ is the fluorescence intensity of the donor/acceptor-labelled 5cHPCS or 5CS/3CS before and after completion of strand displacement reaction kinetics, respectively. *t*_0_ is the start time of the reaction. The fitted pseudo first order reaction rate for strand transfer reactions is *k*_obs’_.

The strand transfer reaction rates, *k*_obs’_, were plotted against the concentration of non-labelled 3CS (Figure 3C) and were observed to linearly vary with increasing concentrations ([3CS)]. As mentioned earlier, the linear relationship between *k*_obs’_ and [3CS] follows equation [2] (27–29). The obtained kinetic parameters for strand transfer reactions are *k*_ass’_ and *k*_diss’_.

Obtained kinetic parameters by fitting the linear plot of *k*_obs’_ against increasing [3CS] with equation [2] are shown in Table 1. Based on the acquired kinetic parameters coupled to the continuous increase in donor intensity plateau (Figure 2F and S3C), a reaction mechanism can be proposed for both DENV2C promoted-3CS displacement from either 5cHPCS/3CS (Scheme 2) or 5CS/3CS (Scheme b in supplementary):

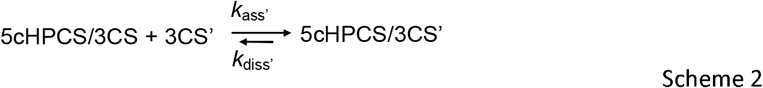

where the final displaced duplex, 5cHPCS/3CS’ or 5CS/3CS’, is formed through a bimolecular reaction (29, 30) and is governed by the second order association constant, *k*_ass’_, and the first order dissociation constant, *k*_diss’_, respectively. The observed values of the *k*_ass’_ and *k*_*diss’*_ were 6.8 (± 0.2) × 10^3^ M^−1^s^−1^ and 14.4 (± 2.5) × 10^−4^ s^−1^, respectively for 5cHPCS/3CS’ strand displacement (Table 1). Interestingly, the comparison of *k*_*ass*_ and *k*_*ass’*_ values indicates that the 5cHPCS/3CS annealing is ~20-folds faster as to that of the 5cHPCS/3CS’ strand displacement. Therefore, the strand displacement reactions/events require the presence of DENV2C in chaperoning the displacement of strands from annealed ORNs complexes, in line with the literature (20–24). The corresponding *k*_ass’_ and *k*_*diss’*_ values for the 5CS/3CS’ strand displacement were found to be 13.3 (± 0.4) × 10^3^ M^−1^s^−1^ and 35.5 (± 5.5) × 10^−4^ s^−1^, respectively (Table 1). Furthermore, the postulated strand displacement mechanisms for both 5cHPCS/3CS’ and 5CS/3CS’ were validated using the Dynafit numerical resolution software (35), by simultaneously fitting the experimental progress curves obtained at different [3CS] (Figure S2). Again, the estimated rate constants *k*_ass’_ and *k*_diss’_ (Table S1) were in excellent agreement with those found by the empirical approach (Table 1). Interestingly, DENV2C-promoted strand displacement of 3CS from the 5cHPCS/3CS annealed complex (5cHPCS/3CS’) was found to be ~2-fold slower as compared to DENV2C-promoted 5CS/3CS’ strand displacement (by comparing the corresponding *k*_*ass’*_ values in Table 1), suggesting that the cHP element plays a structural function in modulating both annealing as well as strand displacement reactions.

### DENV2C-promoted stand displacement of 3CS initiates at the 3’ end of the 5cHPCS sequence

We further characterize the molecular mechanism of 5cHPCS/3CS’ strand displacement by investigating the nucleation of 3CS displacement. To determine whether the 3CS is displaced at the bottom of the cHP stem or at the 3’ end of the 5cHPCS overhang, we used a 32-nt 3cHPCS RNA sequence, where 3CS is elongated by adding complementary nucleotides corresponding to the 5cHP region (Figure 1C). If 3CS displacement starts from the bottom of the cHP stem of the 5cHPCS/3CS annealed duplex, slower kinetic parameters as compared to 5cHPCS/3CS’ strand displacement are expected due to presence of two cHP-like structures which need to be annealed before displacing 3CS. On the other hand, no or minimal changes in the kinetic parameters are expected if the displacement of 3CS from the 5cHPCS/3CS annealed duplex is nucleated from the 3’ end of the 5cHPCS sequence, as no changes in the FAM fluorescence will be observed due to annealing of two cHP-like structures.

The real-time fluorescence intensity traces of DENV2C-promoted 3CS strand displacement from the 5cHPCS/3CS annealed duplex in the presence of increasing concentrations of the 3cHPCS sequence (Figure S4C) were fitted to a monoexponential equation [1]. Further, the reaction rates, *k*_obs’_, were plotted against the concentration of non-labelled 3cHPCS (Figure 3C) and was observed to be linearly varying with increasing concentrations ([3cHPCS)]. The linearly varying kinetic parameters coupled to the continuous increase in donor intensity plateau again suggest that the reaction mechanism is similar to that of 5cHPCS/3CS’ (Scheme c in Supplementary). Interestingly, the observed values of 7.3 (± 0.4) × 10^3^ M^−1^s^−1^ and 12.9 (± 4.5) × 10^−4^ s^−1^ for *k*_ass’_ and *k*_*diss’*_ respectively for 5cHPCS/3cHPCS’ strand displacement are similar to the corresponding values of 5cHPCS/3CS’ (Table 1). Therefore, our results suggest that the displacement of 3CS from the 5cHPCS/3CS annealed duplexes nucleates at the 3’ end of the 5cHPCS.

### DENV2C specifically promotes 3CS displacement

Next, we determined the specificity of 3CS strand displacement from the 5cHPCS/3CS duplex in DENV2C-promoted 5cHPCS/3CS’ strand displacement. For this, we performed strand displacement of the acceptor-labelled 3CS from 5cHPCS/3CS duplex using non-labelled 23-nt 5cHP’, 21-nt 3UAR and 21-nt 5UAR’ sequences (Figure 1C). We compared the efficiency of these three sequences in displacing the 3CS from 5cHPCS/3CS annealed duplex with the native non-labelled 3CS sequence. The 5cHP’ sequence is complementary to the 5cHP hairpin region of 5cHPCS sequence. Therefore, 5cHP’ sequence should not displace acceptor-labelled 3CS due to melting of the cHP hairpin (Figure S5). Furthermore, both 3UAR and 5UAR’ sequences show either non- or partial-complementarity to the 5cHPCS sequences and again should not displace acceptor-labelled 3CS if the 5cHPCS/3CS’ strand displacement is specific (Figure S5). As expected, minimal to no increase in FAM fluorescence was observed upon addition of either 5cHP’, 3UAR or 5UAR’ sequences as compared to the native 3CS sequence (Figure 4A and 4B), suggesting that the specific complementarity between 3CS and 5CS is a requisite for DENV-promoted 5cHPCS/3CS’ strand displacement.

**Figure 4:**
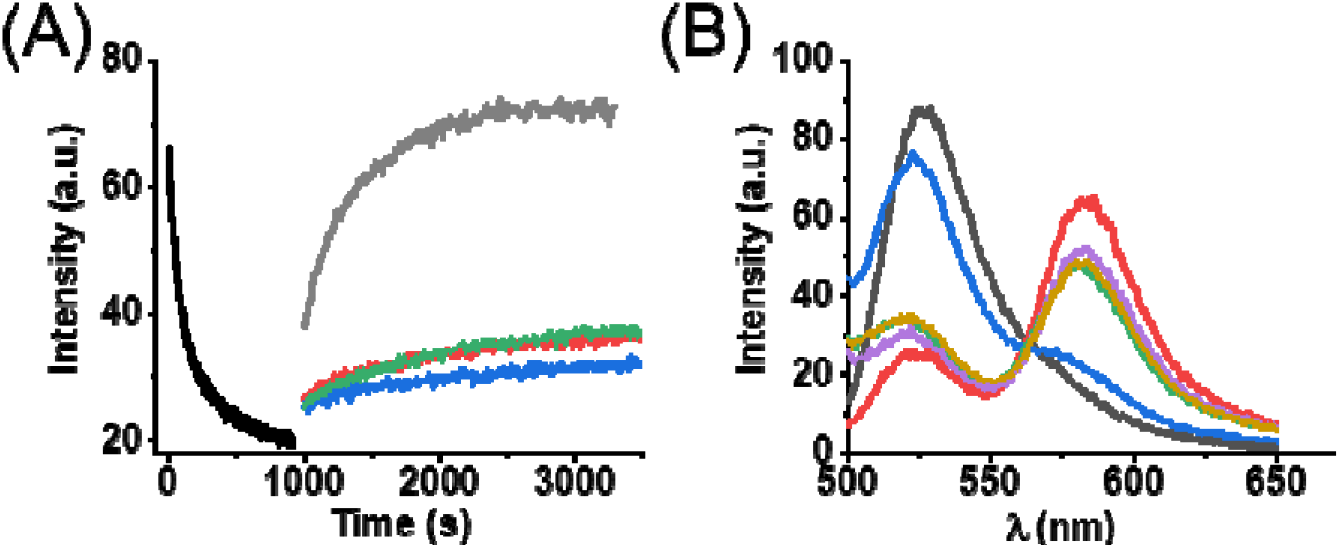
Specificity in the displacement of the 3CS sequence. (A) Real time progress curves of annealing (black trace) between 10nM FAM-labelled 5cHPCS with 10 nM TAMRA-labelled 3CS. Strand displacement of acceptor-labelled 3CS from preformed 5cHPCS/3CS annealing duplex (black trace) in the presence of 300 nM non-labelled 3CS (grey trace), 300 nM 5cHP’ (red trace), 300 nM 3UAR (blue trace) and 300 nM 5UAR’ (green trace). All reactions were performed in the presence of 2 μM DENV2C. The excitation and emission wavelengths were 480 nm and 520 nm, respectively. Fitting annealing curve (black trace) with equation [1] provided *k*_*obs*_ = 3.2 × 10^−3^ s^−1^. Fitting of strand displacement reactions with equation [1] provided values for 5cHPCS/3CS’ (*k* = 3.7 × 10^−3^ s^−1^), 5cHPCS/5cHP’ (*k*_*obs’*_ = 3.4 × 10^−5^ s^−1^), 5cHPCS/3UAR (*k*_*obs’*_ = 3.2 × 10^−5^ s^−1^) and 5cHPCS/5UAR’ (*k*_*obs’*_ = 4.5 × 10^−5^ s^−1^). The 5cHPCS/5cHP’, 5cHPCS/3UAR and 5cHPCS/5UAR’ reactions were 2 orders of magnitude slower as compared to the 5cHPCS/3CS’ reaction, showing the displacement of 3CS from annealed duplex. (B) Emission spectra of 10nM FAM-labelled 5cHPCS before (grey) and after (red) the annealing reaction with 10 nM TAMRA-labelled 3CS. Emission spectra acquired at the end of the strand displacement reactions for 5cHPCS/5cHP’ (pink), 5cHPCS/3UAR (green), 5cHPCS/5UAR’ (yellow) and 5cHPCS/3CS’ (blue). The excitation wavelength was 480 nm.

### DENV2C protein chaperones 5cHPCS/3CS and 5CS/3CS annealing and 5cHPCS/3CS’ and 5CS/3CS’ strand displacement by stabilizing the kinetically active duplex

We have earlier investigated DENV2C chaperone activity during annealing of various essential RNA elements from the 5’UTR (18). Here we focused on understanding how DENV2C chaperones the 5cHPCS/3CS’ and 5CS/3CS’ strand displacement. For this, we investigated if DENV2C favors any kinetically active duplex as is the case of annealing of complementary sequences (18) using trFRET. In trFRET, the energy transfer from donor to acceptor influences the donor fluorescence lifetime, τ, in a distance-dependent manner. The closer the donor and acceptor dyes are, the faster the donor dye relaxes to ground state and the shorter the lifetime. The fluorescence of 5cHPCS had an average lifetime (<τ_avg_>) of 3.8 ± 0.3 ns (Table 2) and was best fitted with two discrete lifetime components of 5.2 ± 0.3 ns (<τ_1_>) and 2.06 ± 0.2 ns (<τ_2_>), having populations of 54 ± 7 % (α_1_) and 46 ± 13 % (α_2_) respectively (Table 2). The existence of more than one population for donor-only labelled 5cHPCS sequence indicates that FAM fluorescence in the cHP hairpin is quenched by neighboring nucleotides probably due to at least two different conformations of the 5CS overhang (Figure S6A). Moreover, we cannot exclude the existence of conformations with very short lifetimes or with lifetimes close to the two principal components, which would not be resolve in our experimental conditions. Similarly, we observed two different conformations for donor-only labelled 5CS sequence with an average lifetime of 3.6 ± 0.6 ns (Table 2) and two discrete lifetime components of 5.13 ± 0.5 ns (<τ_1_>) and 2.09 ± 0.3 ns (<τ_2_>) having populations of 51 ± 4 % (α_1_) and 49 ± 9 % (α_2_), respectively (Table 2).

**Table 2:** Lifetime parameters of 5cHPCS, 5cHPCS/3CS and 5CS/3CS duplexes in the absence and presence of DENV2C. The lifetime traces for single (5cHPCS), dual-labelled duplex (5cHPCS/3CS) were fitted to bi-exponential decay model, and the average fluorescence lifetimes (<τ_avg_>) of FAM for donor were calculated in the absence and in the presence of DENV2C. Fitting of the fluorescence decay provided two discrete lifetimes, <τ_1_> and <τ_2_> with the corresponding fraction of each population as α_1_ and α_2_, respectively. Fitting of the fluorescence decay of FAM in the presence of acceptor (TAMRA) labelled ORN sequences provided two discrete lifetimes, <τ_1_> and <τ_2_’> with the corresponding fraction of each population as α_1_ and α’_2_, respectively. For lifetime traces in the presence of DENV2C, the protein was added at molar concentration of 2 μM, as indicated in material and methods section. Error bars represents standard deviations of at least three repeats.

| ORN | $\langle\tau_{avg}\rangle$ (ns) | $\alpha_1$ (%) | $\alpha_2$ (%) | $\alpha_2'$ (%) | $\langle\tau_1\rangle$ (ns) | $\langle\tau_2\rangle$ (ns) | $\langle\tau_2'\rangle$ (ns) |
| --- | --- | --- | --- | --- | --- | --- | --- |
| 5cHPCS | 3.8 ($\pm 0.3$) | 54 ($\pm 7$) | 46 ($\pm 13$) | | 5.21 ($\pm 0.3$) | 2.06 ($\pm 0.2$) | |
| 5cHPCS/3CS | 3.07 ($\pm 0.4$) | 40 ($\pm 5$) | | 60 ($\pm 7$) | 5.64 ($\pm 0.4$) | | 1.33 ($\pm 0.4$) |
| 5cHPCS/3CS + DENV2C | 2.6 ($\pm 0.2$) | 33 ($\pm 8$) | | 67 ($\pm 6$) | 5.36 ($\pm 0.2$) | | 1.24 ($\pm 0.1$) |
| 5CS | 3.64 ($\pm 0.6$) | 51 ($\pm 4$) | 49 ($\pm 9$) | | 5.13 ($\pm 0.5$) | 2.09 ($\pm 0.3$) | |
| 5CS/3CS | 3.03 ($\pm 0.2$) | 40 ($\pm 10$) | | 60 ($\pm 5$) | 5.57 ($\pm 0.4$) | | 1.32 ($\pm 0.3$) |
| 5CS/3CS + DENV2C | 2.6 ($\pm 0.3$) | 29 ($\pm 10$) | | 71 ($\pm 10$) | 5.32 ($\pm 0.6$) | | 1.40 ($\pm 0.4$) |

With the addition of acceptor-labelled 3CS sequence and by forming the dual-labelled 5cHPCS/3CS duplex, the average lifetime of the 5cHPCS decreased to 3.07 ± 0.4 ns (Table 2). The decay curve was best fitted with two discrete lifetime components of 5.6 ± 0.4 ns (<τ_1_>) and 1.3 ± 0.4 ns (<τ’_2_>) having populations of 40 ± 5 % (α_1_) and 60 ± 13 % (α’_2_), respectively (Table 2). The similarity between the values of <τ_1_> (~5.6 ns) when compared the absence of acceptor-labelled 3CS (~5.2 ns) probably indicates the free form of the 5cHPCS sequence. Consequently, a corresponding decrease in the values of the second discrete lifetime (<τ_2_> = 2.06 ns vs <τ’_2_> = 1.3 ns) suggests the existence of the 5cHPCS sequence in hybridized/annealed form. Their corresponding populations are shown as α_1_ and α’_2_ (Table 2). A similar result was obtained with the addition of acceptor-labelled 3CS to the donor-labelled 5CS sequence, indicating the similarities of the 5CS-3CS duplex (Table 2). Interestingly, an additional 7 to 9 % increase in duplex formation was observed, when annealing of 3CS with 5cHPCS or 5CS occurred in the presence of DENV2C. This increase of duplex formation takes place probably at the expense of conformations that were not annealed. The trFRET results explain that DENV2C probably exerts its RNA chaperone function during both annealing and strand displacement by increasing, as well as stabilizing, kinetically active double stranded 5cHPCS/3CS or 5CS/3CS duplexes. This mechanistic behavior of DENV2C during both these events of DENV genome recombination supports the ‘entropy exchange model’ in which a highly flexible protein undergoes disordered-to-ordered transition upon binding to RNA, that in turn leads to the recombination of the RNA structure through an entropy exchange process (23).

## DISCUSSION

Coding-region elements that are required for RNA synthesis have been characterized in other positive-strand viruses like human rhinovirus-14 and human rhinovirus-2 (HRV-14 and HRV-2) (39–41), poliovirus (42), coxsackievirus B3 (43) and encephalomyocarditis viruses (44). In all viruses, these elements are found to be essential for viral RNA synthesis (40, 43–46). Within the *Flaviviridae* family, Hepatitis C Virus (HCV) features similar internal structure located near the 3’ end of the genome in the NS5B-coding region (47, 48). A number of *cis* elements in the UTRs of flavivirus genomes have been shown to perform multiple roles in the viral life cycle (5, 6, 7, 56–59, 49–55). The cHP is the first element in the flavivirus coding region for which two distinct functions have been characterized: a role in translation start codon selection (13) and a role in RNA synthesis (12). As a replication element, the cHP performs in a sequence-independent manner, which would imply that the overall topology of the 5’ end and selection of the first codon by the scanning ribosome are important for promoting RNA synthesis. Moreover, in promoting RNA synthesis, these replication elements could allow for various conformations of vRNA as sequences anneal and melt. One such conformational large-scale genome rearrangement is the 5’-3’ panhandle formation during (+) RNA cyclization and its melting during vRNA linearization. Therefore, investigating the annealing and strand displacement mechanism of the specific *cis* elements like cHP and CS elements would extract information about the functioning of the cHP region during vRNA synthesis. In this study, we have shown using both annealing and strand displacement kinetics that the presence of the 5cHP hairpin structure regulates both annealing as well as strand displacement of the 3CS sequence to and from the 5CS region. trFRET data showed that 5cHPCS sequence exists in two different conformations (Figure S6A) that may regulate both annealing and strand displacement kinetics. Although we observed that 5cHP modulates annealing of the 5CS sequence with its complementary 3CS sequence, no strand displacement was observed, thus specifying the need of the RNA chaperones. RNA chaperones like DENV2C are vital for preventing RNA misfolding by allowing RNA to escape unfavorable kinetic conformations (18, 20, 31, 32, 60–66). Therefore, we characterized the role of DENV2C by investigating the annealing and strand displacement kinetics of the 5cHPCS and 5CS to their complementary 3CS sequences. Due to its chaperone properties, DENV2C accelerates 5cHPCS/3CS annealing while inducing the 5cHPCS/3CS’ strand displacement reaction. Our results also showed that DENV2C specifically nucleates strand displacement of the 3CS without disturbing the structural functionality of the cHP hairpin. Overall, DENV2C-promoted annealing coupled with strand displacement occurs via a favored active conformation of the 5cHPCS, where the 5CS region is probably unstructured (compare between 5cHPCS and 5cHPCS_1_ species in Figure 5). The population of this conformation increases from ~46% to ~67% in the presence of DENV2C. Since DENV2C does not melt the stem region of the 5cHP hairpin (18), this reactive species of the 5cHPCS probably arises from the melting of C_23_-G_32_ and A_24_-U_31_ base pairs in the 5CS hairpin (Figure 5 and S6B). This DENV2C-promoted reactive species (5cHPCS in Figure 5) probably led to an enhanced annealed product formation from ~16% to ~30% in our working conditions (Figure S6C). In addition, the incremental annealed product formation with increasing concentration of [3CS] led to the observed continuous decrease in the donor intensity plateau (Figure 2B and 2E). Interestingly, the ~2-fold increase in annealed product formation (Figure S6C) correlates well with a similar increase in the DENV2C-promoted kinetic rates (compare *k*_*ass*_ values in table 1). The strand displacement event resulting in an RNA duplex caused by a third, invading RNA molecule relates to the process of RNA annealing. RNA chaperones destabilize double strands, starting from the ends or bulges of the base-paired region, independent of the thermodynamic stability of the double strand (67). A third strand can utilize such destabilized regions as starting points for invasion (Figure 5). The concerted process of opening the initial double strand and annealing the new duplex finally results in either the replacement of the original strand (3CS sequence) or the expulsion of the invading strand (3CS’ sequence), according to the kinetics and thermodynamic conditions. We observed that DENV2C could catalyze the strand displacement reaction either by destabilizing edges or by favoring the annealing reaction of the invading strand. Therefore, the proposed mechanism for DENV2C-promoted 3CS strand displacement from the 5cHPCS/3CS annealing duplex propagates through the formation of an intermediate triplex (see 5cHPCS/3CS-3CS’ complex in Figure 5). This intermediate triplex originates due nucleation of the 3CS strand displacement from the 5cHPCS/3CS annealed duplex via 3’ end of the 5cHPCS (Figure 5) and subsequent displacement of acceptor-labelled 3CS sequence. However, the complete displacement of the 3CS from the 5cHPCS/3CS annealed duplex probably takes place only at ~200-fold excess of displacing sequences ([3CS’] and [3cHPCS]) (Figure S6D). Similarly, a reaction mechanism is proposed for DENV2C-promoted coupled annealing and strand displacement of 5CS with its complementary 3CS sequences (Figure S7).

**Figure 5:**
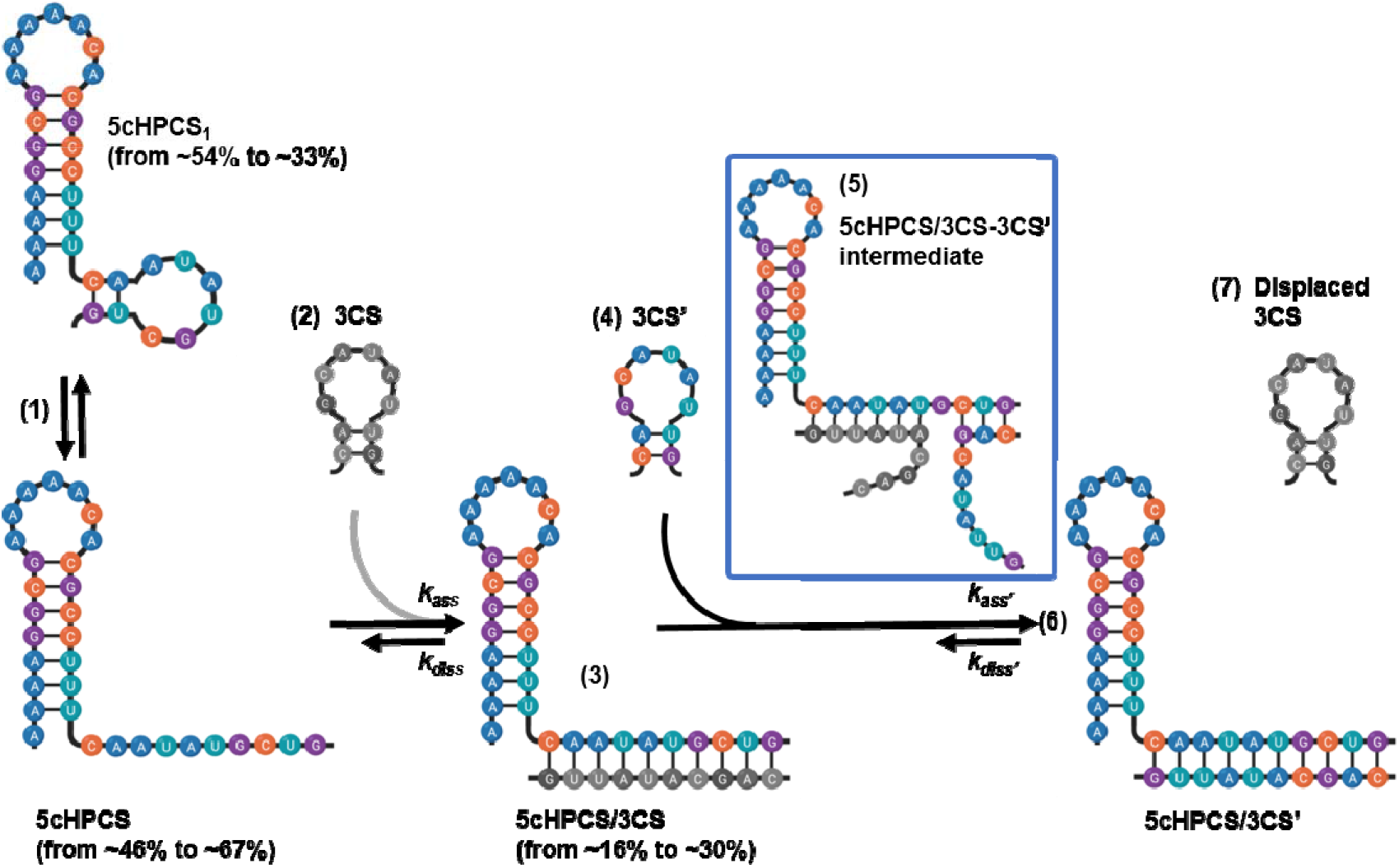
Proposed reaction mechanisms for 5cHPCS/3CS annealing and 5cHPCS/3CS’ strand displacement in the presence of DENV2C. The proposed reaction mechanism shows an equilibrium (step 1) between at least two different species of the 5cHPCS (5cHPCS and 5cHPCS_1_) that originates due to opening of the 5CS hairpin through melting C_23_-G_32_ and A_24_-U_31_ base pairs. This melting of base pairs increases from 46% to 67% in the presence of DENV2C that in turn accelerates the annealing reaction in the presence of the complementary 3CS sequence (step 2) and leads to higher (up from 16% to 30%) and rapid (2-fold faster) annealed duplex formation (step 3). DENV2C catalyzes the strand displacement reaction in the presence of at least 10-fold molar excess of displacing strand 3CS’ (step 4) by destabilizing the 5CS-3CS duplex at the 3’ end of 5cHPCS sequence. This destabilizing of the 3’ edge of the 5CS-3CS duplex lead to nucleation of a transient intermediate triplex (step 5), 5cHPCS/3CS-3CS’, that originates from 3’ end of the 5cHPCS (shown in the blue color rectangle). The strand displacement reaction leads to the formation of an annealing duplex with 3CS’ (step 6) due to the displacement of 3CS (step 7).

The mechanism by which DENV2C rearranges the viral genome during RNA synthesis is one of the most obscure steps of the DENV life cycle. Here, we provide mechanistic insights in which the structure of cHP could stabilize 5’-3’ panhandle and how DENV2C modulates the function of the cHP during the 5CS-3CS annealing and strand displacement. To our knowledge, this is the first demonstration of DENV2C chaperoning strand displacement from an annealed duplex. These abilities of DENV2C are critical for genomic RNA dimerization in DENV replication and RNA packaging as well as in facilitating recombination between various DENV genotypes and subtypes to increase viral variability. Moreover, delineation of RNA structures in the presence of RNA chaperones like DENV2C serve to broaden our understanding of the mechanisms viruses use to reproduce and to subvert or evade the host antiviral response, and allows for the focused development of vaccines and antiviral therapies.

