## Supplementary Materials for "Dengue virus 2 capsid protein chaperones strand displacement without altering the capsid-coding region hairpin element’s structural functionality"

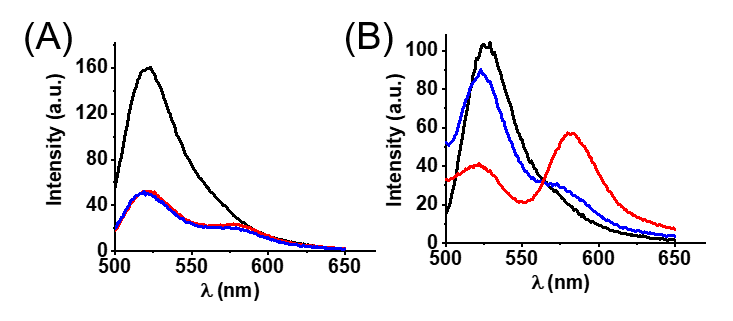

**Figure S1: Emission spectra of 5cHPCS before (black), after annealing with 3CS (red) and after strand displacement with 3CS’ (blue) in the (A) absence and (B) presence of DENV2C.** (A) Emission spectra before and after annealing of 10 nM 5cHPCS to 10 nM 3CS, and the subsequent displacement of 3CS from the 5cHPCS/3CS duplex. (B) Emission spectra before and after annealing of 10 nM 5cHPCS to 10 nM 3CS, and the subsequent displacement of 3CS from the 5cHPCS/3CS duplex in the presence of DENV2C. The excitation wavelength was 480 nm.

**Dynafit simulation procedure**

Simulated progress curves were obtained at different concentration of 3CS (5 nM, 10 nM, 30 nM, 50 nM, 0.1 μM, 0.2 μM, 0.5 μM and 1 μM) for annealing (Figure S2A) and 3CS’ (0.1 μM, 0.3 μM, 0.5 μM, 1 μM, 1.5 μM, 2 μM, 3 μM and 5 μM) for strand displacement (Figure S2C) reactions using elementary rate constants, *k*_ass_ and *k*_diss_ from Table 1, and according to the following schemes:

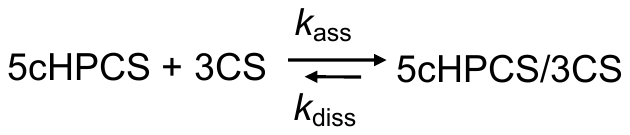
 for annealing

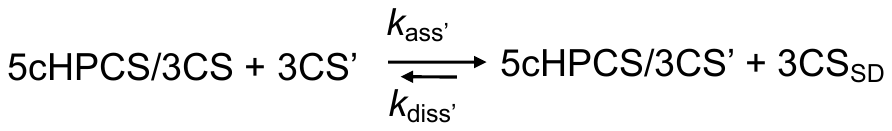
for strand displacement

To obtain simulated progress curves, the concentration of 5cHPCS kept at a constant value of 10 nM. Progress curves of the 5cHPCS/3CS annealing reaction were fitted to equation [1] to obtain values for *k*_obs_ with increasing concentration of 3CS (Figure S2B) in absence (Figure S2B: solid squares) and presence (Figure S2B: solid circles) of DENV2C. Similarly, 5cHPCS/3CS’, 5CS/3CS’ and 5cHPCS/3cHPCS strand displacement reactions were fitted to equation [1] to obtain *k*_obs_*_’_* values (Figure S2D in the presence of DENV2C. The reaction rates, *k*_obs_ and *k*_obs’,_ were plotted against the concentration of complementary/strand displacement ORNs (Figure S2B and Figure S2D) and fitted using equation [2]. The generated kinetic parameters are shown in Table S1.

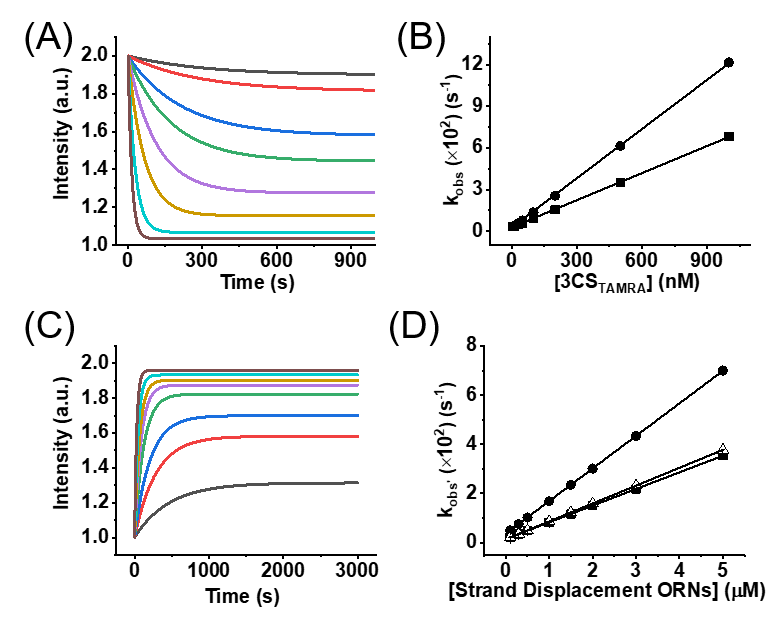

**Figure S2: Simulated kinetic parameters of 5cHPCS/3CS annealing and 5cHPCS/3CS’, 5CS/3CS’ and 5cHPCS/3cHPCS strand displacement.** Simulated progress curves at different concentration of 3CS (5 nM - black, 10 nM - red, 30 nM - blue, 50 nM - green, 0.1 μM - pink, 0.2 μM - yellow, 0.5 μM - cyan and 1 μM - olive) for (A) 5HPCS/3CS annealing in the absence of DENV2C and 3CS’ (0.1 μM - black, 0.3 μM - red, 0.5 μM - blue, 1 μM - green, 1.5 μM - pink, 2 μM - yellow, 3 μM - cyan and 5 μM - olive) for (C) 5cHPCS/3CS’ strand displacement were simulated using scheme 1 and 2, respectively. The obtained *k*_obs_ values for both (B) annealing of 5cHPCS/3CS in the absence (squares) and presence (circles) of DENV2C, and (D) strand displacement of 5cHPCS/3CS’ (squares), 5CS/3CS’ (circles) and 5cHPCS/3cHPCS (open triangles) were plotted against increasing concentrations of complementary/strand displacing ORNs. The solid lines correspond to the fit of the *k*_obs_ values with equation [2]. The values are provided in Table S1. Similar values of simulated kinetic parameters (Table S1) to that of experimental kinetic parameter (Table 1) values support the proposed reaction schemes in the article.

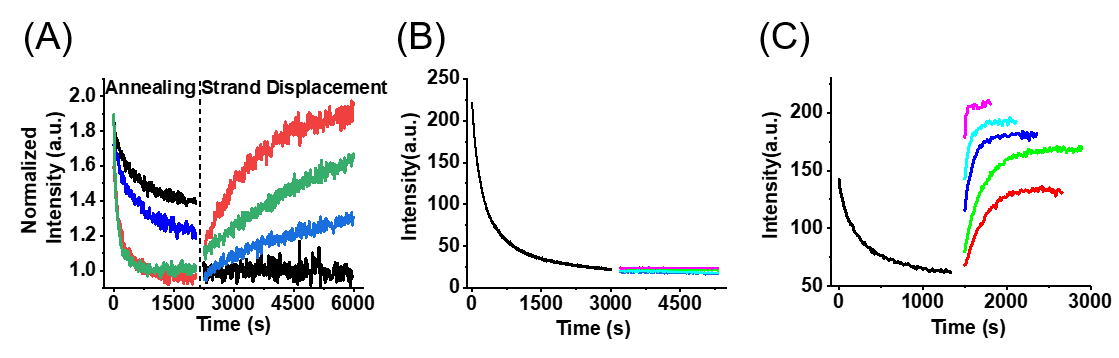

**Figure S3: Dose response plot for different concentrations of DENV2C and progress curves of 5CS/3CS’ strand displacement.** (A) Dose-response plot for 5cHPCS/3CS annealing was performed using 10 nM FAM 5cHPCS and 10 nM TAMRA-labelled 3CS. The subsequent 5cHPCS/3CS’ strand displacement with 300 nM non-labelled 3CS’ was done in the absence (black traces) or in the presence of DENV2C at concentrations of 100 nM (blue traces), 1 μM (green traces) and 2 μM (red traces). Fitting of annealing traces with equation [1] provided the following *k*_obs_ values: 1.8 × 10^-3^ s^-1^ (no DENV2C), 2.3 × 10^-3^ s^-1^ (100 nM DENV2C), 3.4 × 10^-3^ s^-1^ (1 μM DENV2C) and 3.3 × 10^-3^ s^-1^ (2 μM DENV2C). The subsequent *k*_obs’_ values for 5cHPCS/3CS’ strand displacement reactions by fitting with equation [1] are: 9.7 × 10^-5^ s^-1^ (100 nM DENV2C), 2.4 × 10^-4^ s^-1^ (1 μM DENV2C) and 4.4 × 10^-3^ s^-1^ (2 μM DENV2C). No strand displacement reaction was observed in the absence of DENV2C. Observed annealing rates were saturated at 1 μM DENV2C but not the strand displacement rates, and thus DENV2C was used at 2 μM concentration. In addition, at higher DENV2C concentration (>2 μM), reliable and reproducible data could not be extracted due to the rapid nature of both annealing as well as strand displacement reactions. Therefore, 2 μM DENV2C was chosen as the protein concentration to conduct all annealing and strand displacement reactions, unless otherwise stated. **Real-time progress curves of 5CS/3CS annealing and 5CS/3CS’ strand displacement in the absence (B) and presence (C) of DENV2C.** Progress curves of 10 nM FAM-labelled 5CS annealing (black traces) with 10 nM TAMRA-labelled 3CS and 5CS/3CS’ strand displacement initiating from the duplex formed during the 5CS/3CS annealing in the (B) absence (red-100 nM, green-500 nM, blue-1 μM, cyan-1.5 μM and pink-2 μM) and in the (C) presence (red-100 nM, green-500 nM, blue-1 μM, cyan-1.5 μM and pink-2 μM) of DENV2C. All annealing and strand transfer traces were fitted using equation [1] and the obtained values are plotted in Figure 3. Excitation and emission wavelengths used were 480 nm and 520 nm, respectively.

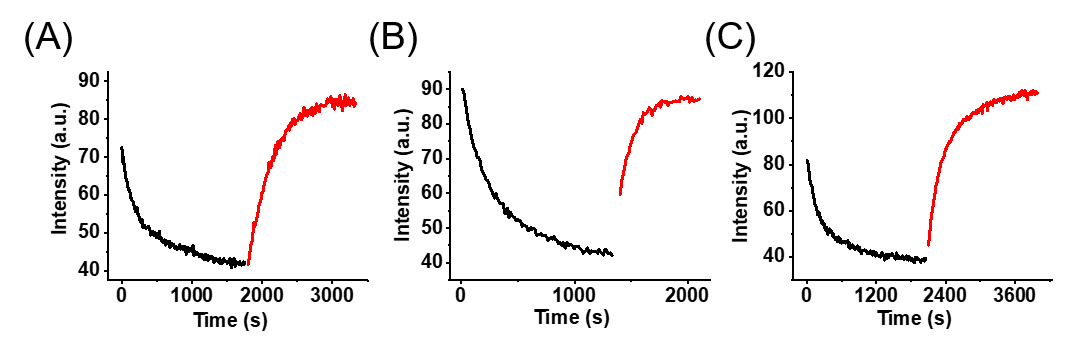

**Figure S4: Comparison of real-time progress curves of annealing and strand displacement reactions in the presence of DENV2C.** Progress curves of 10 nM FAM-labelled (A and C) 5cHPCS and (B) 5CS annealing (black traces) with 10 nM TAMRA-labelled 3CS. The fitting of annealing curves with equation [1] provided values of *k*_obs_ = 3.2 × 10^-3^ s^-1^ and *k*_obs_ = 4.8 × 10^-3^ s^-1^ for 5cHPCS/3CS and 5CS/3CS annealing, respectively. Excitation and emission wavelengths used were 480 nm and 520 nm respectively. Strand displacement progress curves (red traces) in the presence of (A) 300 nM 3CS’, (B) 300 nM 3CS’ and (C) 300 nM 3cHPCS from preformed (A and C) 5cHPCS/3CS and (B) 5CS/3CS annealed duplexes, respectively. The fitting of strand displacement curves with equation [1] provided values of *k*_obs’_ = 3.7 × 10^-3^ s^-1^, *k*_obs’_ = 8 × 10^-3^ s^-1^ and *k*_obs’_ = 3.3 × 10^-3^ s^-1^ for 5cHPCS/3CS’, 5CS/3CS’ and 5cHPCS/3cHPCS strand displacement, respectively. Excitation and emission wavelengths used were 480 nm and 520 nm respectively.

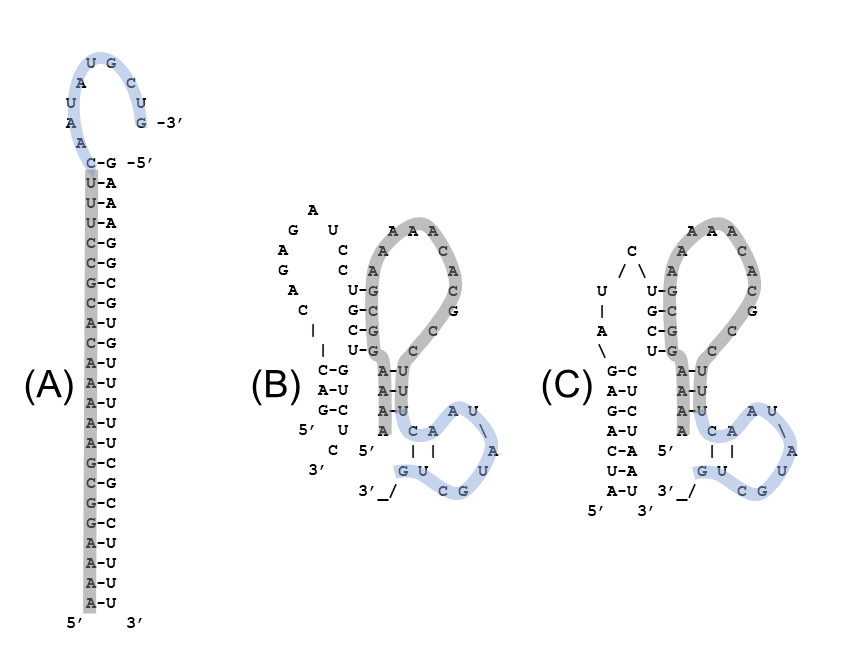

**Figure S5: Extended duplex structures.** Extended duplexes of (A) 5cHPCS/5cHP’, (B) 5cHPCS/3UAR and (C) 5cHPCS/5UAR’. To show the region specific hybridization in the 5cHPCS sequence, the 5cHP sequence is highlighted in grey while the 5CS region segment is highlighted in blue. A dash indicates complementary base pairing. The duplex structures were determined using the Two-state melting (hybridization) webtool from http://unafold.rna.albany.edu/. The duplex formation shows no to minimum complementary hybridization of 5cHP’, 3UAR and 5UAR’ with the 5CS region of the 5cHPCS.

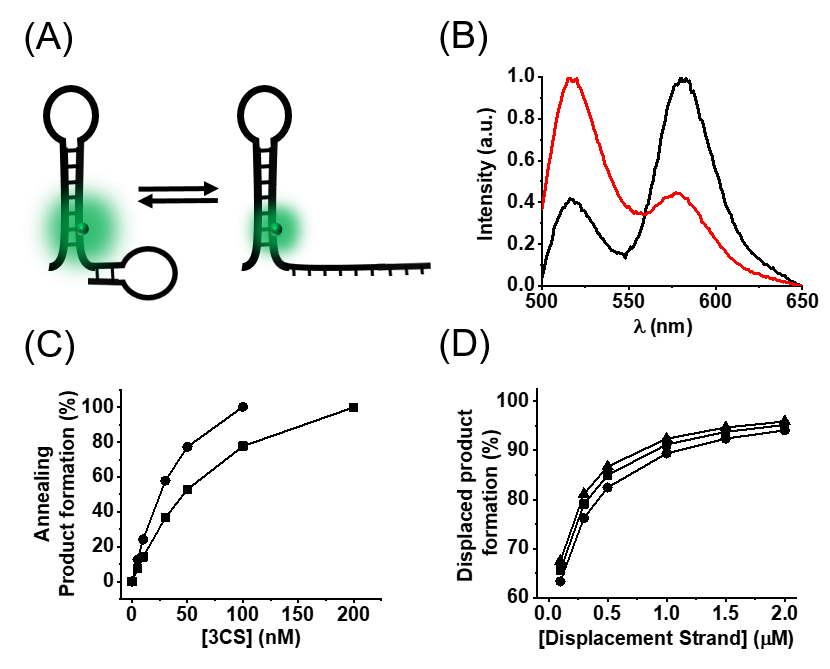

**Figure S6: Different 5cHPCS conformations during annealing and strand transfer product formation. (A)** Schematic representation of two possible conformations of the 5cHPCS sequence. The open conformation originates due to the melting of C_24_-G_32_ and A_25_-U_31_ base pairs in the 5CS region of the 5cHPCS. The melting of base pairs probably leads to the unhindered movement of the 5CS overhang in 5cHPCS and in turn lowers the donor intensity possibly, due to its quenching with the colliding 5CS nucleotides. **(B)** We tested the melting of C_24_-G_32_ and A_25_-U_31_ base pairs using doubly-labelled 5CS with FAM as the donor fluorophore at the 5’ end and TAMRA as the acceptor fluorophore at the 3’ end, forming a Förster resonance energy transfer (FRET)-pair. Due to the proximity of donor and acceptor dyes in hairpin conformation of the 5CS, the fluorescence of FAM is quenched (black). With the addition of DENV2C, the hairpin converts into a single-stranded sequence due to melting of base pairs in the stem region, in turn increasing the distance between donor and acceptor fluorophores. This increase in distance leads to florescence recovery of the donor (red). Percentages of **(C)** annealing product and **(D)** strand displaced product were calculated using simulated data shown in Figure S2. Product formation was calculated using the intensity plateau for both annealing as well as strand displacement reaction at different concentrations of complementary/displacement strands as against highest concentration used in annealing (1 μM) and strand displacement (5 μM) reactions. **(C)** For 5cHPCS/3CS annealing the percentages were calculated in absence (circles) and presence (squares) of DENV2C. **(D)** Similarly for strand displacement, the percentages were calculated for 5cHPCS/3CS’ (squares), 5CS/3CS’ (circles) and 5cHPCS/3cHPCS (triangles) in the presence of DENV2C.

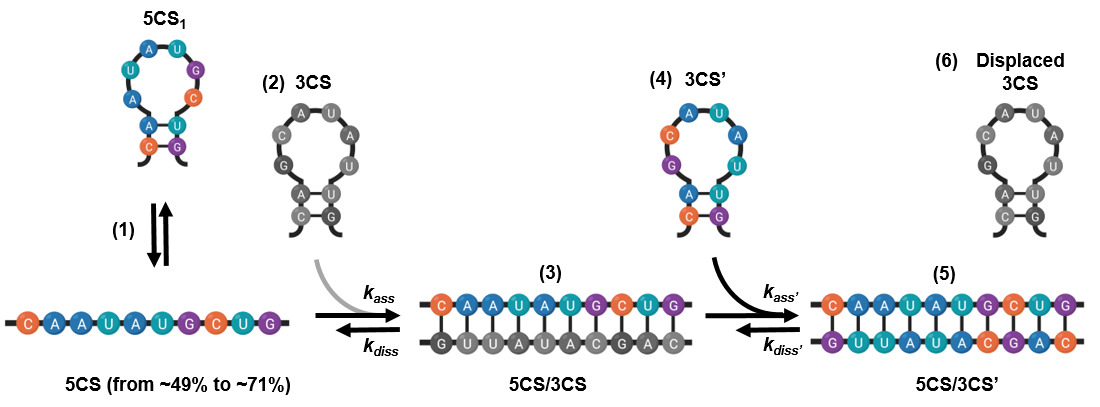

**Figure S7: Proposed reaction mechanisms for 5CS/3CS annealing and 5CS/3CS’ strand displacement in the presence DENV2C.** The proposed reaction mechanism shows an equilibrium (step 1) between at least two different species of the 5CS (5CS and 5CS_1_) that originates from the opening of the 5CS hairpin through the melting of C_1_-G_10_ and A_2_-U_9_ base pairs. This melting of base pairs increases from 49% to 71% in the presence of DENV2C, which in turn accelerates the annealing reaction (step 2) in the presence of the complementary 3CS sequence (step 3). DENV2C catalyze the strand displacement reaction in the presence of at least 10-fold molar excess of displacing strand 3CS’ (step 4) by destabilizing the 5CS-3CS duplex probably at the 3’ end of 5CS sequence. The strand displacement reaction led to the formation of the annealing duplex with 3CS’ (step 5) due to the displacment of 3CS (step 6).

**Table S1:** **Kinetic parameters of 5cHPCS/3CS and 5CS/3CS annealing, and 5cHPCS/3CS’, 5CS/3CS’ and 5cHPCS/3cHPCS strand displacement in the absence and presence of DENV2C obtained by fitting simulated progression curves.**

|  | | | **Annealing** | |  | **Strand Displacement** | |
| --- | --- | --- | --- | --- | --- | --- | --- |
| **Fluorescein labelled ORN** | **TAMRA-labelled Complementary ORN** | **DENV2C**  **(μM)** | ***k*_ass_**  **(M^-1^s^-1^) × 10^-3^** | ***k*_diss_**  **(s^-1^) × 10^4^** | **Non-labelled Strand Displacement ORN** | ***k*_ass’_**  **(M^-1^s^-1^) × 10^-3^** | ***k*_diss’_**  **(s^-1^) × 10^4^** |
| **5cHPCS** | **3CS** | 0 | 65.1 (±0.5) | 28.6 (±0.5) | **3CS’** | - | - |
| **5CS** | **3CS** | 0 | 101.6 (±0.5) | 41.4 (±1.2) | **3CS’** | - | - |
| **5cHPCS** | **3CS** | 2 | 118.3 (±0.8) | 26.7 (±1.4) | **3CS’** | 6.8 (±0.05) | 14.7 (±0.1) |
|  |  |  |  |  | **3cHPCS** | 7.3 (±0.06) | 13.2 (±0.1) |
| **5CS** | **3CS** | 2 | 194.7 (±1) | 33.4 (±1.3) | **3CS’** | 13.3 (±0.1) | 36.1 (±0.1) |

Kinetic rate constants were calculated from the dependence of the *k*_obs_ values on the concentration of the labelled/unlabelled complementary ORNs. The *k*_ass_, *k*_diss_, *k*_ass’_ and *k*_diss’_ values were calculated using equation [2]. Simulated kinetic parameters and empirical values (Table 1) were similar, supporting the proposed reaction schemes in the article.

**Reaction schemes:**

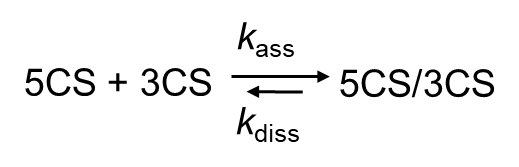
 Scheme a

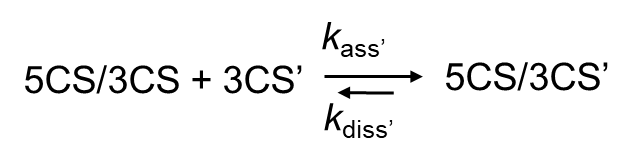
 Scheme b

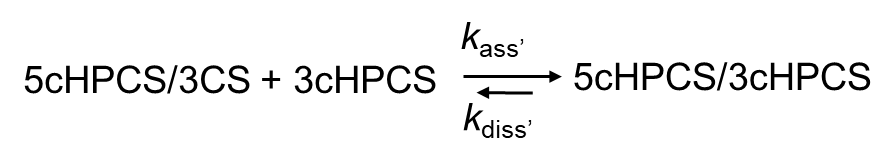
 Scheme c

According to the above-mentioned reaction schemes, a reaction mechanism with a single kinetic pathway is proposed.
